# Pre-existing immunity provides a barrier to airborne transmission of influenza viruses

**DOI:** 10.1101/2020.06.15.103747

**Authors:** Valerie Le Sage, Jennifer E. Jones, Karen A. Kormuth, William J. Fitzsimmons, Eric Nturibi, Gabriella H. Padovani, Claudia P. Arevalo, Andrea J. French, Annika J. Avery, Richard Manivanh, Elizabeth E. McGrady, Amar R. Bhagwat, Adam S. Lauring, Scott E. Hensley, Seema S. Lakdawala

## Abstract

Human-to-human transmission of influenza viruses is a serious public health threat, yet the precise role of immunity from previous infections on the susceptibility to airborne viruses is still unknown. Using human seasonal influenza viruses in a ferret model, we examined the roles of exposure duration and heterosubtypic immunity on influenza transmission. We found that airborne transmission of seasonal influenza strains is abrogated in recipient animals with pre-existing non-neutralizing immunity, indicating that transmissibility of a given influenza virus strain should be examined in the context of ferrets that are not immunologically naïve.

## Introduction

Airborne transmission is essential for the epidemiological success of human influenza A virus (IAV), which imposes a significant seasonal public health burden. Every influenza season is different, with one virus subtype (H3N2 or H1N1) typically dominating and factors such as age, pregnancy and pre-existing medical conditions putting people at increased risk of severe influenza infection. During the 2017-2018 H3N2 predominant season, roughly 79,000 people died in the United States, which is more than the number who died during the 2009 H1N1 pandemic (*1, 2*). In the 2017-2018 H3N2 epidemic, 40% of the cases were in the elderly (65+), in contrast during the 2009 H1N1 pandemic the highest burden of infection was found in individuals 5-24 years of age (48%) (*1, 2*). This age-based discrepancy in IAV burden suggests that pre-existing immunity could impact the susceptibility to IAV infection, since people of different age groups are exposed to different strains of IAV in early childhood (*3–6*).

An individual’s first influenza infection typically occurs before the age of 5 (*7*) and are repeatedly infected with influenza viruses during their lifetime. The antibody response to a person’s first influenza infection will be boosted upon each subsequent antigenically distinct influenza virus strain in a process classically referred to as ‘Original Antigenic Sin’ (*8*). The impact of pre-existing immunity on the spread of influenza viruses has been understudied, and the few reports in this area have found that pre-existing immunity against heterologous or homologous strains surprisingly protects animals against all subsequent influenza virus infections (*9, 10*). In these studies, Steel et al demonstrated that pre-existing seasonal H1N1 or H3N2 immunity in recipient guinea pigs reduced transmission of the 2009 H1N1 pandemic virus (*10*) and Houser et al observed that pre-existing seasonal H3N2 immunity in donor ferrets prevented transmission of emerging 2011 swine H3N2v virus (*9*). However, these observations do not mimic human epidemiology, since individuals can become infected with influenza virus many times, suggesting that these animal models may need to be revised to more accurately represent human transmission.

The majority of published transmission studies use 14 days of continuous exposure in immune naïve animals (*11–14*) and a time interval of 4-6 weeks between primary and secondary infection (*9, 10*). To address these transmission parameters, we examine the role of timing of exposure and pre-existing immunity to address the barriers to transmission and provide a comprehensive comparison of a seasonal H3N2 virus and the 2009 pandemic H1N1 virus (H1N1pdm09). We used ferrets for respiratory droplet transmission because they are naturally susceptible to human isolates of IAV and can transmit infectious IAV particles through the air (*12, 15, 16*). We propose that transmissibility of emerging influenza viruses be assessed in immune ferrets and at short exposure times. In addition, our results indicate that pre-existing immunity provides different barriers for H3N2 and H1N1pdm09 virus transmission, suggesting that pre-existing immunity can drive susceptibility to heterosubtypic infections.

## Results

### IAV transmission to naive animals is efficient after short and periodic exposures

To investigate the constraint of exposure time on transmission, we examined a representative human seasonal H3N2 virus (A/Perth/16/2009) and the most recent IAV pandemic H1N1pdm09 virus (A/CA/07/2009). Airborne transmission of these two viruses to naïve ferrets was performed continuously for 7 or 2 days, as well as periodically for 8 hours a day for 5 consecutive days (Fig 1A). These times were used to mimic human exposure conditions. H1N1pdm09 transmitted by respiratory droplets to 100% of all naïve recipients at all exposure times. Donor ferrets shed from days 1 to 5 post-infection (Fig 1B-D, red bars), while recipient ferrets had a wider range of shedding (Fig 1B-D, blue bars). Shorter exposure times caused a delay in detectable H1N1pdm09 virus in recipient nasal secretions consistently after day 4 post-exposure (Fig 1C and D, blue bars). This observation is consistent with the highly transmissible nature of this virus in multiple transmission systems (*14, 17*). The H3N2 virus replicated efficiently in donor ferrets on days 1,2, 3 and 5 (Fig 1E-G, green bars). H3N2 transmitted to 3/3 naïve animals after a 7-day exposure time with slightly delayed shedding kinetics on days 2 to 7 post-infection (Fig 1E, green bars), as compared to ferrets infected with H1N1pdm09 (Fig 1B, red bars). After a 48 h exposure, H3N2 transmitted to 4/4 naïve recipients (Fig 1F) and 2/3 in a second independent replicate (Fig S1), with recipient animals shedding on days 4, 6 and 8 post-infection (Fig 1F and S1, orange bars). At intermittent exposure times transmission of H3N2 virus was slightly reduced to 2/3 naïve recipients, which we still consider to be efficient airborne transmission. Taken together, these results indicate that seasonal influenza viruses are highly transmissible to naïve recipients.

**Figure 1.**
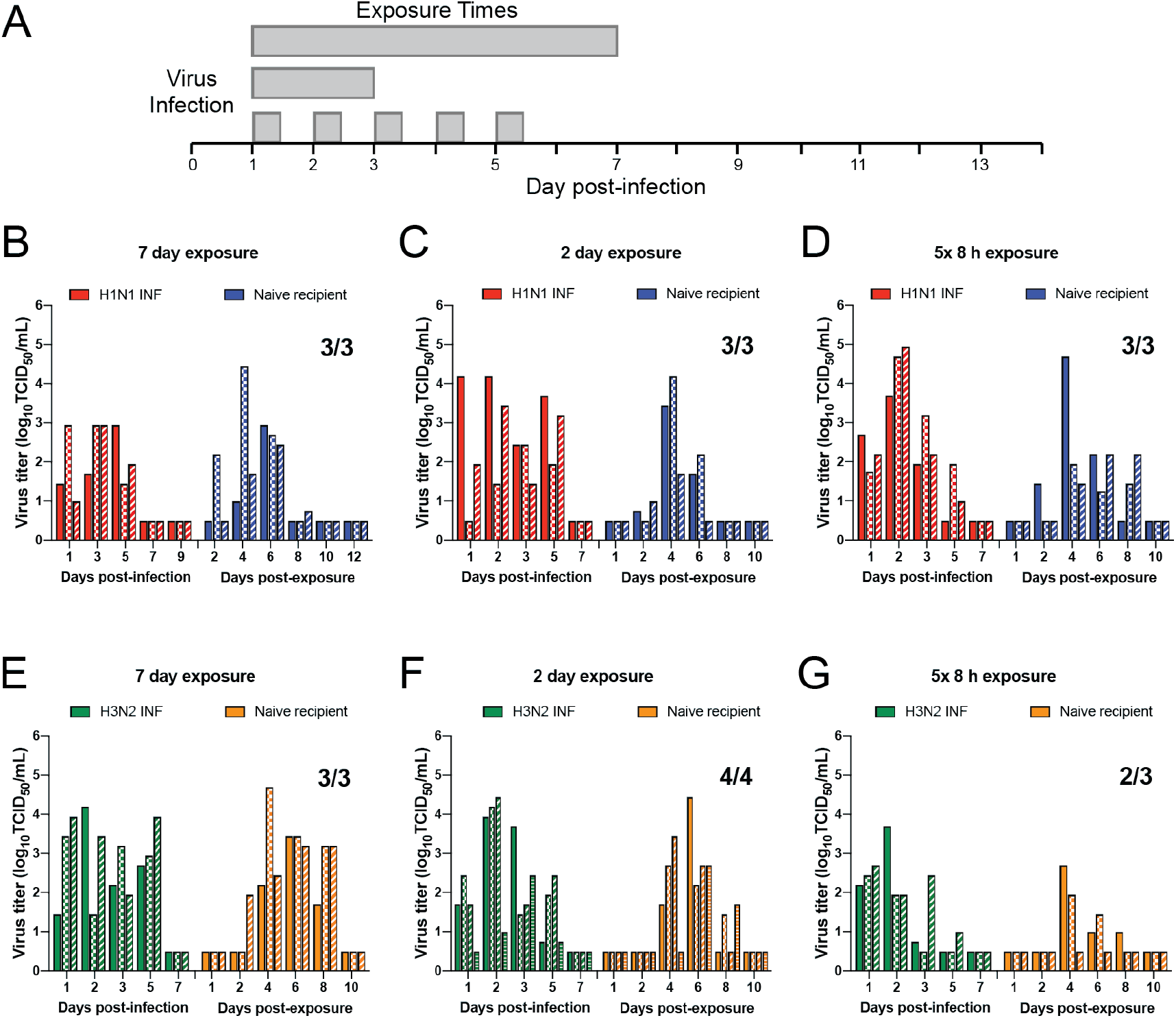
Efficient transmission of seasonal H1N1 and H3N2 influenza viruses to naive animals under short or periodic exposures. (A) Schematic of experimental procedure. Shaded gray bars depict exposure times. Three donor ferrets were infected with A/CA/07/2009 (H1N1) (B-D) or A/Perth/16/2009 (H3N2) (E-G) and a naïve recipient ferret was placed in the adjacent cage at 24 hour post-infection for 7 (B and E) or 2 (C and F) continuous days, or 8 hours a day for 5 consecutive days (D and G). Nasal washes were collected from all ferrets on the indicated days and each bar indicates an individual ferret. Limit of detection is indicated by dashed line. Viral titers for donor animals in part G have previously been published in (*18*).

### Pre-existing immunity to heterologous strains impacts airborne transmission of influenza viruses at short exposure times

As demonstrated in Figure 1, ferrets are naturally susceptible to human H1N1pdm09 and H3N2 influenza virus infections. To develop a model to mimic the yearly seasonality of influenza infections, we waited between 60 and 84 days between infections to allow for the primary immune response to wane, as shown by others (*19–22*), and result in two robust infections. Six ferrets were infected with H3N2 virus and then three of those animals were experimentally infected at 60 days post-infection with H1N1pdm09 virus (herein referred to as ‘H3-H1 INF’) (Fig 2A). Twenty-four hours after this secondary infection, the three remaining ferrets with H3 pre-existing immunity (Table S1) were each placed in an adjacent cage to act as the recipient animal (herein referred to as ‘H3-imm recipients’) and exposed for 2 days (Fig 2A). An 84-day gap produced a robust infection of the H1N1pdm09 virus in experimentally inoculated H3-H1 INF ferrets with virus detected in ferret nasal secretions on multiple days (Fig 2B). We observed that transmission of H1N1pdm09 to recipient animals was not significantly impacted by pre-existing H3N2 virus immunity as 2/3 H3-imm recipients became infected after a 2-day exposure period (Fig 2B). Interestingly, the duration and kinetics of H1N1pdm09 shedding were different between naïve animals and H3-imm animals (Fig 1C vs Fig 2B).

**Figure 2.**
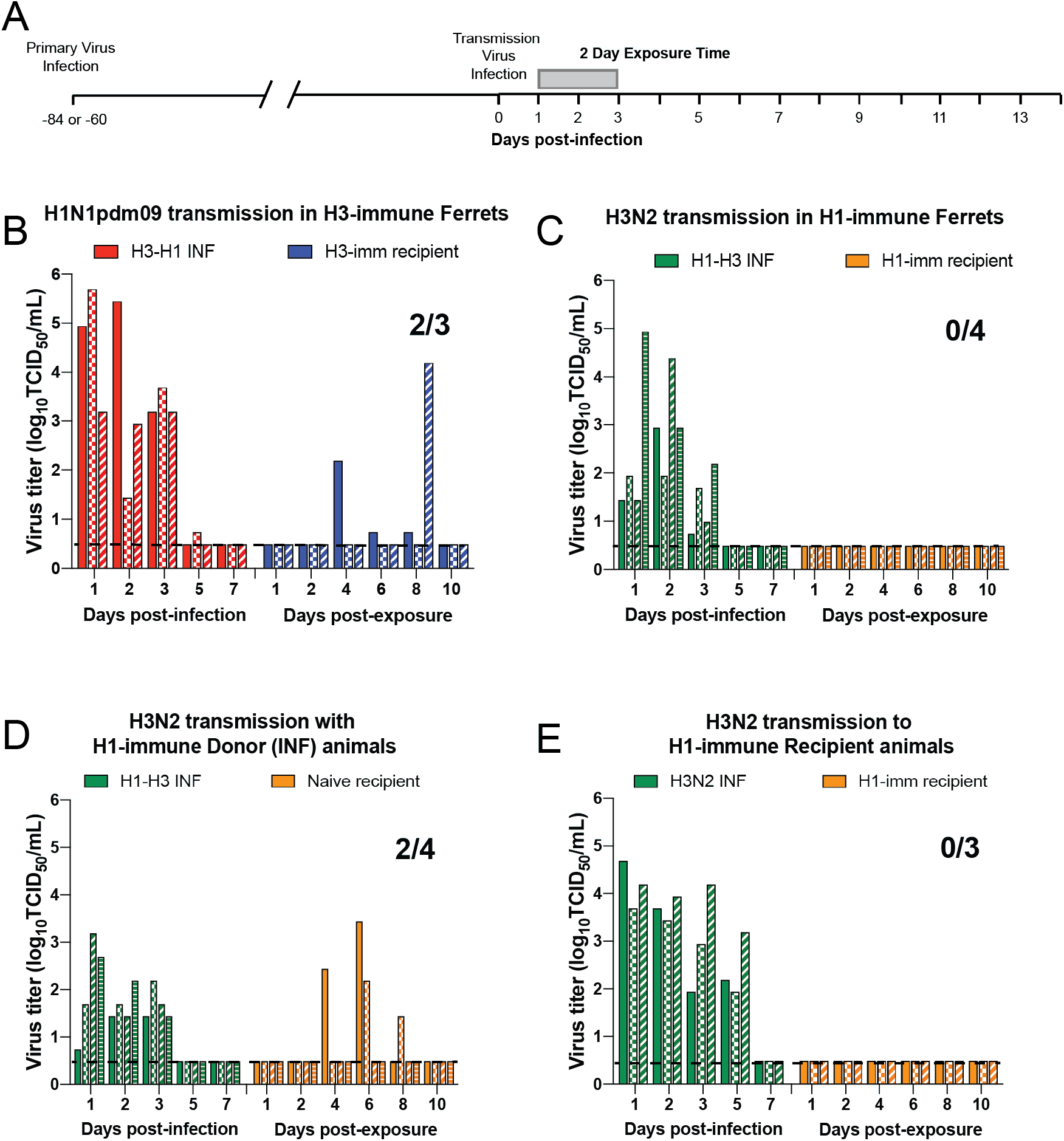
Pre-existing heterosubtypic immunity blocks H3N2 transmission but not H1N1pdm09 transmission. (A) Schematic of experimental procedure, gray bars depict 2-day exposure time. (B) All donor and recipient animals were infected with A/Perth/16/2009 (H3N2) virus 60 days prior to H1N1pdm09 (A/CA/07/2009) transmission. Donor animals are denoted as H3-H1 INF, and recipient animals are ‘H3-imm’. (C) All donor and recipient animals were infected with A/CA/07/2009 (H1N1pdm09) virus 84 days prior to H3N2 (Perth/16/2009) transmission. Donor animals are denoted as H1-H3 INF, and recipient animals are ‘H1-imm’. The contribution of donor and recipient immunity was tested separately for A/Perth/16/2009 (H3N2) transmission. Donors only (D) or recipient only (E) had pre-existing immunity to A/CA/07/09 H1N1pdm09. Donor animals are denoted as H1-H3 INF or H3N2 INF (no prior immunity), and recipient animals are either ‘naïve’ to indicate no prior immunity or ‘H1-imm’. Each bar represents an individual animal. Limit of detection is denoted by a dashed line.

In a complementary study, ferrets with pre-existing H1N1pdm09 immunity were experimentally infected with H3N2 virus to act as donors (herein referred to as ‘H1-H3 INF’) in a subsequent transmission experiment 84 days later (Fig 2A). Robust replication of H3N2 in H1N1pdm09 immune donor (H1-H3 INF) animals was observed (Fig 2C, green bars) and a recipient ferret with pre-existing H1N1pdm09 immunity (herein referred to as ‘H1-imm recipient’) was placed in the adjacent cage 24 h post-infection. The recipient animals were exposed to the H1-H3 INF donor animals for 2 continuous days. Surprisingly, no virus was detected in the nasal secretions of the H1-imm recipients (Fig 2C, orange bars) and no seroconversion was observed on day 13 postexposure (Table S1). We previously demonstrated that H3N2 was able to transmit to 6 out of 7 recipients with no prior immunity (Figure 1F and S1). In comparison to this efficient transmission, pre-existing H1N1pdm09 immunity provides a complete block to H3N2 transmission during a 2-day exposure window (Fig 2C).

To discern whether donor or recipient immunity was critical for the blockade in H3N2 virus transmission, we examined the spread of H3N2 virus to either H1-imm donors or recipients. In Figure 2D, H1-H3 INF donors transmitted the virus to 2/4 recipients without prior immunity (50%) (Fig 2D, orange bars). This H3N2 transmission efficiency is reduced as compared to 85% (6/7) in animals without prior immunity (Fig 1F and S1), indicating that donor immunity may partially contribute to a barrier in H3N2 transmission. In contrast, 0/4 H1-imm recipients became infected upon a 2-day exposure to ferrets infected with H3N2 virus as their primary infection (Fig 2E, orange bars), which suggests that heterosubtypic immunity in recipients is sufficient to block H3N2 virus transmission. Taken together, these results indicate that airborne transmission of H3N2 cannot overcome the barrier imposed by H1N1pdm09 immunity but transmission of H1N1pdm09 can overcome H3N2 immunity even within short exposure times.

### Barrier to H3N2 airborne transmission is independent of neutralizing antibodies

To elucidate the difference in protection provided by H1N1pdm09 versus H3N2 immunity, we first compared the shedding of the naïve and immune donor ferrets. Higher viral loads in the donor animals might be expected to result in more efficient transmission, yet they do not always correlate with each other (*23*). Direct comparison of the shedding kinetics between transmission experiments was not possible as the H1N1pdm09 and H3N2 nasal washes were titered at different times on different cell lines. However, comparison of viral RNA levels in nasal washes of H1 or H3 infected animals revealed similar levels between these two viral infections during both primary and secondary infections (Fig S2A-B). These results show that the block in H3N2 transmission was not due to significantly decreased viral shedding by ferrets with pre-existing H1N1pdm09 immunity.

To determine whether the H1-imm recipient ferrets were generating cross-protective H3 neutralizing antibodies, we performed neutralization assays with sera from animals after primary or secondary infection. Sera from the H3-imm and H3-H1 INF animals exhibited robust H3N2 neutralizing antibodies that persisted from day 14 post infection/exposure and appeared to increase slightly upon heterologous challenge (Fig 3A). Sera from the H3-imm recipient animals, had no detectable cross-reacting neutralizing antibodies against H1N1pdm09 (Fig 3B) with the H1N1pdm09 virus transmitting efficiently to 2/3 H3-imm recipient animals (Fig 2B). Similarly, sera from H1-imm ferrets produced strong H1N1pdm09 neutralizing antibody titers that waned slightly over 84 days and increased slightly upon the H3 transmission experiment (Fig 3D). Although the H1-imm recipient animals were protected against H3 transmission they had no detectable crossreacting neutralizing antibodies against H3 (Fig 3C, orange line). At day 97 post-infection, only the experimentally infected animals (H1-H3 INF) had neutralizing antibody titers against H3 (Fig 3C, green line). These data indicate that the block in H3N2 airborne transmission by H1N1pdm09 immunity is independent of cross-reactive neutralizing antibodies in either the donor or recipient.

**Figure 3.**
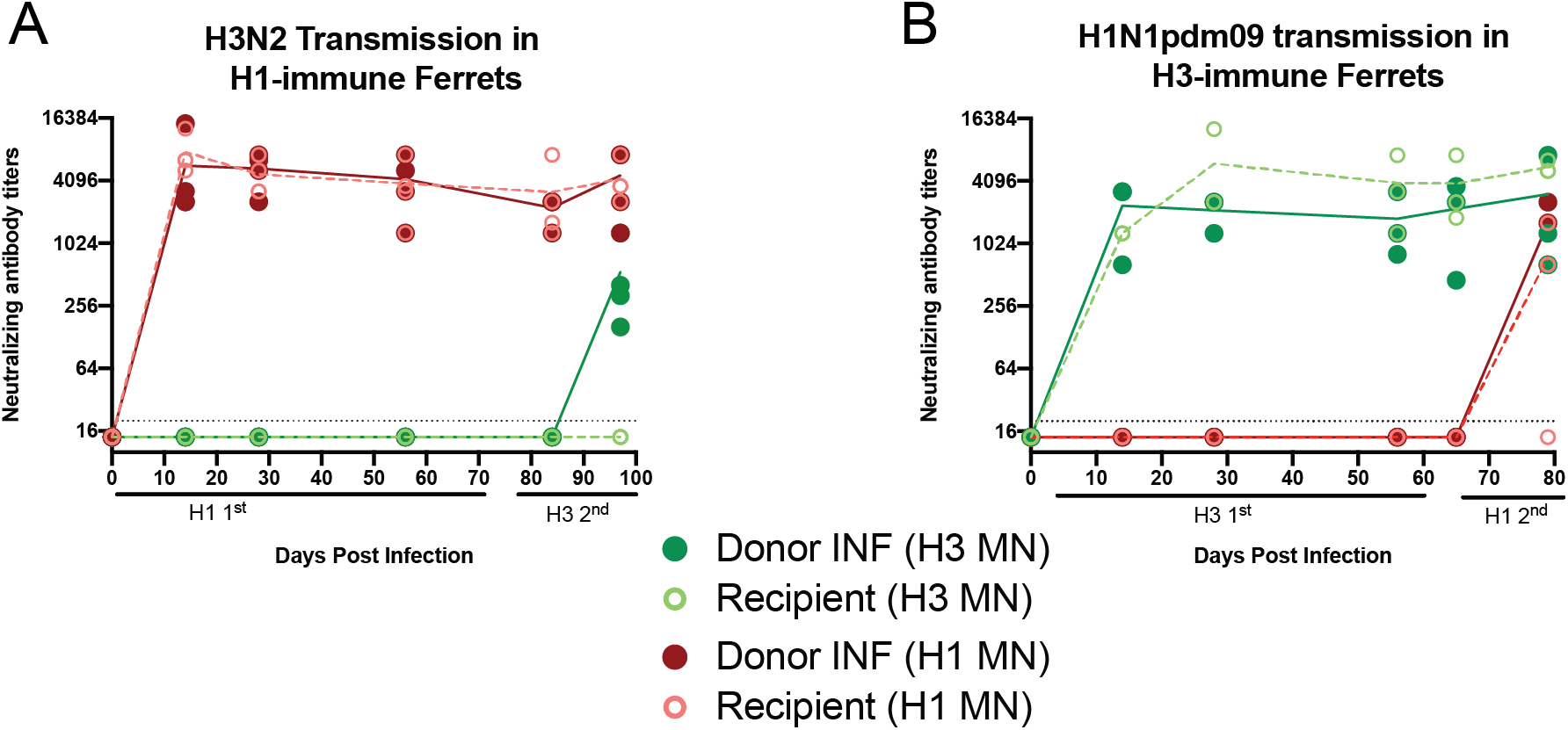
Neutralizing antibody titers against H1N1 and H3N2 viruses. We examined the neutralizing antibody titers against A/CA/07/09 (H1N1pdm09; red lines) and A/Perth/16/09 (H3N2; green lines) from ferrets depicted in Fig 2B and C. Sera from animals from Fig 2C; H1N1pdm09 primary infection followed by H3N2 transmission are depicted in panel A. Sera from animals from Fig 2B, H3N2 primary infection followed by H1N1pdm09 transmission, are depicted in panel B. At the indicated times, serum was collected and neutralizing antibodies against both H1N1pdm09 (red lines) and H3N2 (green lines) were determined by microneutralization assay. Donor infected animals are denoted in solid circles and recipient animals are denoted by open circles. Each point is an individual animals and trend lines for donor (solid) or recipient (dashed) animals are shown. Limit of detection is indicated by the dashed line.

## Discussion

Airborne transmission is critical for emergence of pandemic viruses and we demonstrate that epidemiologically successful human seasonal and pandemic influenza viruses transmit to naïve recipients efficiently even at shortened or intermittent exposure times. In humans, primary influenza infections typically occur by about 5 years of age, which initiates innate and adaptive immune responses resulting in immunological memory. Thus, the spread of pandemic and seasonal IAV occurs in a population with significant pre-existing immunity (*23*). We demonstrate that pre-existing immunity can influence airborne transmissibility of IAV. Specifically, transmission of seasonal H3N2 virus is abrogated in animals with pre-existing H1N1pdm09 immunity that was non-neutralizing. Conversely, H3N2 immunity did not significantly impair H1N1pdm09 transmission. Importantly, recipient immunity is sufficient to get a complete block in H3N2 transmission through an immune mechanism that does not rely on cross-neutralizing antibodies.

Transmissibility of a given influenza virus relies on the viral replication in the upper respiratory tract, release of virus into the air, survival of the virus within the environment, and successful replication in a susceptible recipient. In the ferret animal model, transmission studies typically house a naïve recipient animal adjacent to an infected donor for 14 consecutive days (*11–14*) and very few studies have altered this experimental animal model. In this study, we examined the transmissibility of seasonal H3N2 and H1N1pdm09 viruses at shortened exposures times and found efficient transmission after a 48 h continuous exposure and intermittent exposure of 8 h a day for 5 days, mimicking a work week. Studies examining transmission of H1N1pdm09 have demonstrated transmission to naïve ferrets after a 30 h exposure to as short as a 3 h exposure (*23, 24*), which is consistent with our observations with H1N1pdm09. However, it is important to note that the transmission system utilized by Koster et al (*23*) had considerably lower air flow rates, 6 liters per minute compared to 40 cubic ft. per minute (Fig S3). Interestingly, transmissibility of H1N1pdm09 within a 30 h exposure window was efficient if the naïve recipients were exposed to infected ferrets early (1-2 days) as opposed to late (5-6 days) post-infection (*25*), suggesting that transmission efficiency is also linked to an exposure window.

The primary correlate of protection against influenza virus infections has been mapped to antibodies against of the two antigenic determinants on the surface of the virus particle, glycoproteins hemagglutinin (HA) and neuraminidase (NA) (*26*). In this study, we also demonstrate that heterosubtypic immunity can provide protection from transmission of H3N2 virus. Broadly protective immune response can be generated between HA subtypes (*3, 4, 27–32*), but the lack of detectable H3N2 neutralizing antibodies in the serum of donor and recipient animals suggests that these broadly cross-reactive neutralizing antibodies are not the source of protection. Sub-neutralizing concentrations of antibodies that recognize the stalk portion of HA have been shown to limit influenza disease through antibody-dependent cell mediated cytotoxicity (ADCC) (*33–36*). Alternatively, non-neutralizing antibodies against the influenza virus conserved antigens, NP, M1 and M2 might be exerting a protective function through ADCC, antibody-dependent phagocytosis or antibody-mediated complement-dependent cytotoxicity (*37*). Heterosubtypic immunity in the recipients provided a robust barrier against infection with seasonal H3N2, but not H1N1pdm09. This difference may be due to the immunogenicity of internal H1N1pdm09 antigens in ferrets compared to H3N2 virus. Future examination of cross-reactive adaptive immune responses between H1N1pdm09 and H3N2 will further our understanding of influenza virus correlates of protection. Finally, heterosubtypic immunity in experimentally infected animals limited disease severity, but not viral shedding in nasal secretions, which may represent a system to study asymptomatic transmission.

In the last 10 years, two pandemic viruses have emerged in the human population; the 2009 pandemic H1N1 virus and SARS-CoV-2. Both of the viruses are efficiently transmitted from person-to-person with expelled aerosols. During the 2009 H1N1 influenza pandemic over 12,000 individuals died of the infection in the United States, with 77% of those deaths were in patients 18-64 year-old (*38*). Recent analyses correlating birth year to imprinted virus would suggest that a large proportion of those individuals would have been imprinted with a seasonal H3N2 virus (*3*) and our data demonstrate the pre-existing immunity against H3N2 virus was not a significant barrier to natural infection of H1N1pdm09 virus. In contrast, in 2017-2018 a drifted H3N2 predominated for which the vaccine was not efficacious, but individuals aged 5-24 had the lowest number of infections in that year (*2*) and would include a large number of individuals imprinted with H1N1pdm09 virus. Previous studies in ferrets have also demonstrated that pre-existing H1N1pdm09 immunity 31 days prior allowed for rapid clearance of an antigenically distinct swine influenza virus and demonstrated reduced disease severity compared to naïve ferrets (*39*). Our data support this idea as immune imprinting with H1N1pdm09 is superior to H3N2 imprinting to protect from infection with heterologous viruses. Although the immunological mechanism underlying this phenotype will require further studies, translation of these results to the current COVID-19 pandemic may be important to understand age-based distributions of SARS-CoV-2 disease severity and susceptibility.

## Acknowledgements

This work was supported by the National Institute of Allergy and Infectious Diseases (HHSN272201400007C, SSL; 1R01AI139063-01A1, SSL; 1R01AI113047, SEH; 1P01AI108686, SEH; CEIRS HHSN272201400005C, SEH). Additional funding for SSL includes the American Lung Association Biomedical Research grant, and a New Initiative Award from the Charles E. Kaufman Foundation, a supporting organization of The Pittsburgh Foundation. ASL is supported by Burroughs Wellcome Fund PATH award.

## Materials and Methods

### Cells and Viruses

MDCK (Madin Darby canine kidney, obtained from ATCC) and MDCK SIAT cells (kind gift from Dr. Stacy Schultz-cherry at St. Jude) were grown at 37 °C in 5 % CO2 in MEM medium (Sigma) containing 5 % Fetal Bovine Serum (FBS, HyClone), penicillin/streptomycin and L-glutamine. Reverse genetics plasmids of A/Perth/16/2009 and A/California/07/2009 were a generous gift from Dr. Jesse Bloom (Fred Hutch Cancer Research Center, Seattle) and were rescued as previously described in (*40*). The viral titers were determined by tissue culture infectious dose 50 (TCID50) using the endpoint titration method on MDCK cells for H1N1pdm09 and MDCK SIAT cells for H3N2 (*41*).

### Animals

Ferret transmission experiments were conducted at the University of Pittsburgh in compliance with the guidelines of the Institutional Animal Care and Use Committee. Five to six month old male ferrets were purchased from Triple F Farms (Sayre, PA, USA). All ferrets were screened for antibodies against circulating influenza A and B viruses by hemagglutinin inhibition assay, as described in (*40*), using the following antigens obtained through the International Reagent Resource, Influenza Division, WHO Collaborating Center for Surveillance, Epidemiology and Control of Influenza, Centers for Disease Control and Prevention, Atlanta, GA, USA: 2018-2019 WHO Antigen, Influenza A(H3) Control Antigen (A/Singapore/INFIMH-16-0019/2016), BPL-Inactivated, FR-1606; 2014-2015 WHO Antigen, Influenza A(H1N1)pdm09 Control Antigen (A/California/07/2009 NYMC X-179A), BPL-Inactivated, FR-1184; 2018-2019 WHO Antigen, Influenza B Control Antigen, Victoria Lineage (B/Colorado/06/2017), BPL-Inactivated, FR-1607; 2015-2016 WHO Antigen, Influenza B Control Antigen, Yamagata Lineage (B/Phuket/3073/2013), BPL-Inactivated, FR-1403.

### Transmission studies

Our transmission caging setup is a modified Allentown ferret and rabbit bioisolator cage similar to those used in (*16, 40*). Additional details on the caging setup can be found in Fig S3. For each study, three to four ferrets were anesthetized by isofluorane and inoculated intranasally with 10^6^ TCID_50_/500uL of A/Perth/16/2009 or A/California/07/2009, they function as the donor or (INF) animals. Twenty-four hours later, a recipient ferret was placed into the cage but separated from the donor animal by two staggered perforated metal plates welded together one inch apart. Nasal washes were collected from each donor and recipient every other day for 14 days. To prevent accidental contact or fomite transmission by investigators, the recipient ferret was handled first and extensive cleaning of all chambers, biosafety cabinet, and temperature monitoring wands was performed between each recipient and donor animal and between each pair of animals which also included glove and anesthesia chamber changes. Sera was collected from donor and recipient ferrets upon completion of experiment to confirm seroconversion. Environmental conditions were monitored daily and ranged between 20-22 °C with 44-50 % relative humidity. To ensure no accidental exposure during husbandry procedures, recipient animal sections of the cage were cleaned first then then infected side, three people participated in each husbandry event to ensure that a clean pair of hands handled bedding and food changes. One cage was done at a time and a 10 min wait time to remove contaminated air was observed before moving to the next cage. New scrappers, gloves, and sleeve covers were used on subsequent cage cleaning.

Clinical symptoms such as weight loss and temperature were recorded during each nasal wash procedure and other symptoms such as sneezing, coughing, lethargy or nasal discharge were noted during any handling events. Animals were given A/D diet twice a day to entice eating once they reached 10% weight loss. A summary of clinical symptoms for each study are provided in Supplemental Table 1.

### RNA Harvest and Quantification of Viral Load

Viral RNA was extracted from ferret nasal washes by processing 200 μl through the Purelink Pro 96 Viral RNA/DNA Purification Kit (Thermofisher 12280-096A). The viral load in each sample was measured by RT-qPCR using primers/probe specific for the open reading frame of segment 7 (M1/M2) of influenza A virus: forward primer 5’-GACCRATCCTGTCACCTCTGAC-3’, reverse primer 5’-AGGGCATTYTGGACAAAKCGTCTA-3’, and probe 5’-(FAM)-TGCAGTCCTCGCTCACTGGGCACG-(BHQ1)-3’. Each reaction contained 5.4 μL of nuclease-free water, 0.5 μL of each primer at 40 μM, 0.1 μL of ROX dye, 0.5 μL SuperScript III RT/Platinum Taq enzyme mix, 0.5 μL of 10 μM probe, 12.5 μL of 2x PCR buffer master mix, and 5 μL of extracted viral RNA. The PCR master mix was thawed and stored at 4°C, 24 h before reaction set-up. The RT-qPCR was performed on a 7500 Fast real-time PCR system (Applied Biosystems) with a machine protocol of 50°C-30min, 95°C-2min followed by 45 cycles of 95°C-15sec, 55°C-30sec. To relate genome copy number to Ct value, we used a standard curve based on serial dilutions of a plasmid control, run in triplicate on the same plate. H3N2 samples were compared to a plasmid containing the M segment of A/Perth/16/2009. H1N1 samples were compared to a plasmid containing the M segment of A/California/07/2009.

### Serology assay

Analysis of neutralizing antibodies from ferret sera was performed as previously described (*40*). Briefly, the microneutralization assay was performed using 10^3.3^ TCID50 of either H3N2 or H1N1pdm09 virus incubated with 2-fold serial dilutions of heat-inactivated ferret sera. The neutralizing titer was defined as the reciprocal of the highest dilution of serum required to completely neutralize infectivity of 10^3.3^ TCID_50_ of virus on MDCK cells. The concentration of antibody required to neutralize 100 TCID50 of virus was calculated based on the neutralizing titer dilution divided by the initial dilution factor, multiplied by the antibody concentration.

## Supplemental Figures

**Figure S1.**
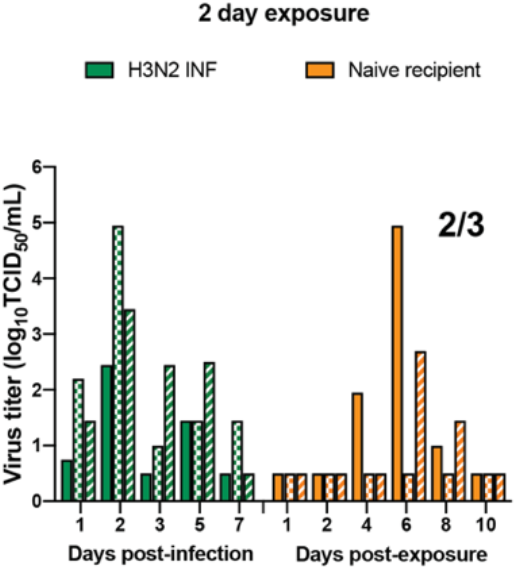
Transmission of H3N2 to naïve ferrets for a short exposure period. Three ferrets were infected with A/Perth/16/2009 (H3N2) and nasal washes were collected from each ferret on the indicated days post-infection. A naive ferret was placed in the adjacent cage at 24 hour post-infection for 2 days and nasal washes were collected from each recipient ferret on the indicated days post-exposure. Bars indicate individual ferrets. All ferrets were serologically negative for circulating influenza viruses at the beginning of the study. The limit of detection was 10^0.5^ TCID_50_/mL. TCID_50_, 50% tissue culture infectious dose. The viral titer data for 3 donor animals was previously published in (*18*).

**Figure S2.**
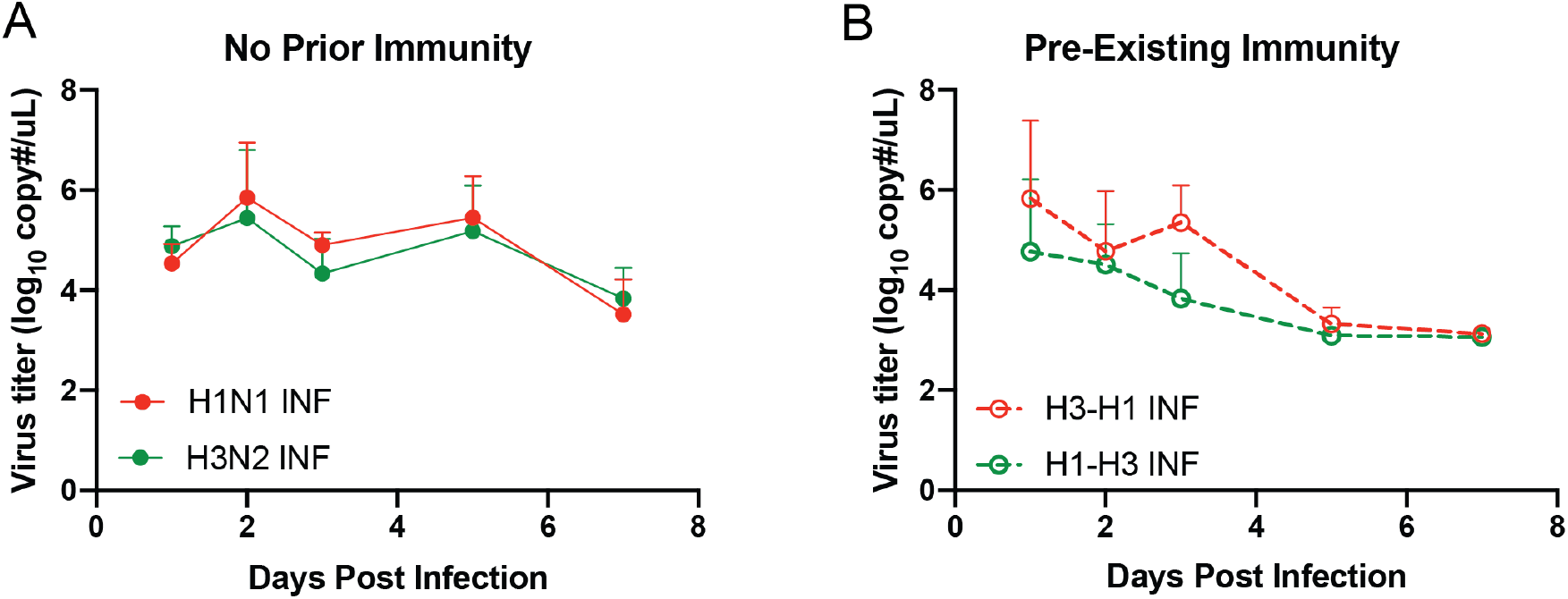
Quantification of H1N1 and H3N2 viral RNA in donor animals. RNA was isolated from nasal wash samples at each of the indicated days post-infection from ferrets infected with H1N1pdm09 (red line) and H3N2 (green line). Data are shown as mean +/- SEM.

**Figure S3.**
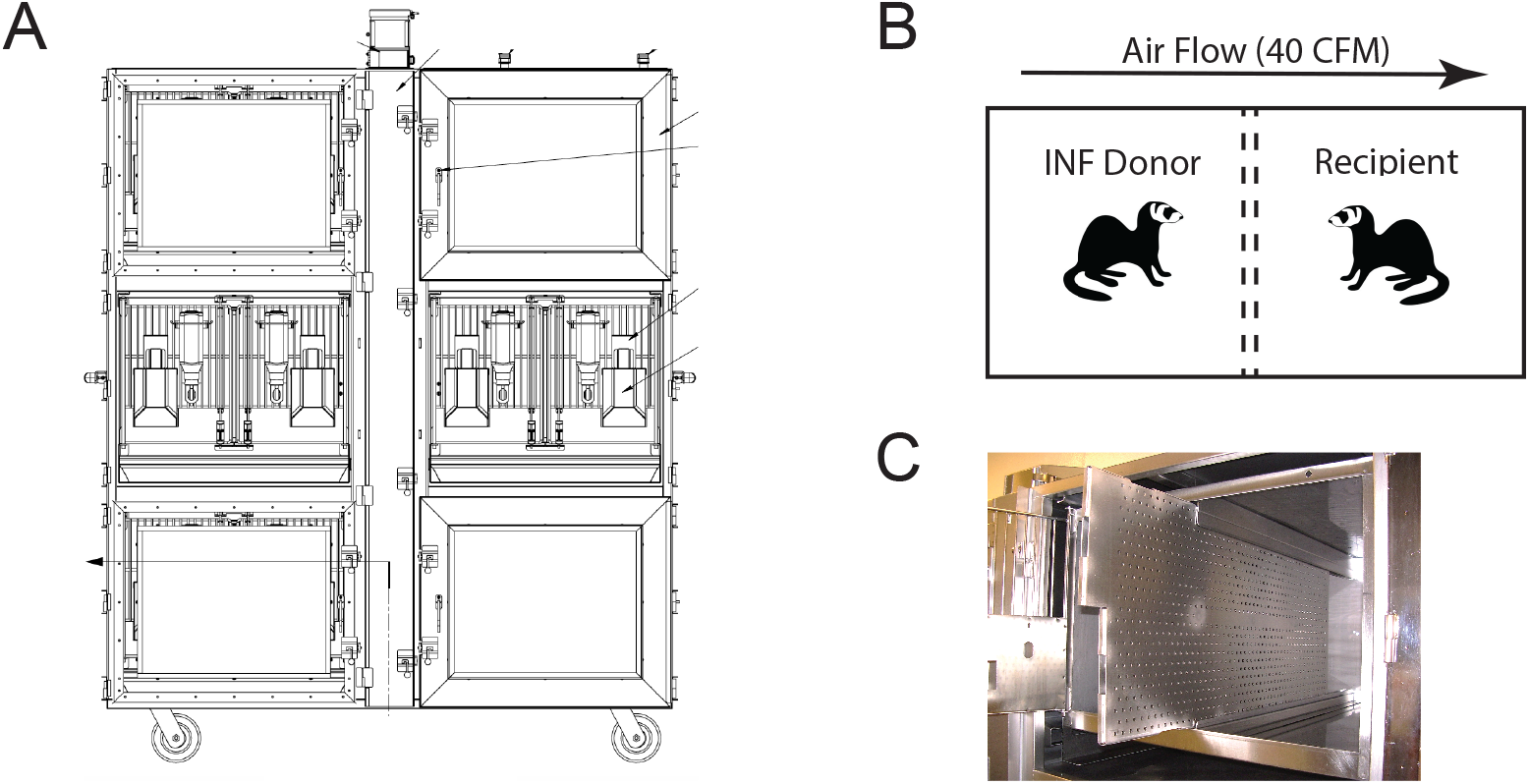
Transmission cage setup. A. Diagram of ferret transmission unit. B. Schematic of ferret experimental setup with constant air flow passing from infected (INF) donor to recipient ferret at 40 cubic feet per minute (CFM). INF donor and recipient are separated by a stainless steel divider, which is made up of two perforated plates, 1 inch apart and staggered holes (5 mm diameter).

**Table S1.** Clinical signs and symptoms.

| Figure | Virus | Exposure time | Status | Transmission efficiency | Temperature increase* | Weight loss* | Sneezing | H1N1 microneutralization titers<sup>^</sup> | H3N2 microneutralization titers<sup>^</sup> |
| --- | --- | --- | --- | --- | --- | --- | --- | --- | --- |
| 1B | H1N1pdm | 7 days | INF |  | 1/3 | 2/3 | 3/3 | 2032, 1280, 1613 | ND |
|  |  |  | Naïve | 3/3 | 2/3 | 2/3 | 3/3 | 1810, 6451, 2560 | ND |
| 1C | H1N1pdm | 2 days | INF |  | 2/3 | 1/3 | 2/3 | 3620, 1280, 1280 | ND |
|  |  |  | Naïve | 3/3 | 1/3 | 1/3 | 2/3 | 1280, 7241, 1280 | ND |
| 1D | H1N1pdm | 5x 8h | INF |  | 2/3 | 0/3 | 1/3 | 14482, 7241, 5120 | ND |
|  |  |  | Naïve | 3/3 | 2/3 | 1/3 | 3/3 | 6451, 5120, 6451 | ND |
| 1E | H3N2 | 7 days | INF |  | 0/3 | 0/3 | 0/3 | ND | 3620, 1810, 1810 |
|  |  |  | Naïve | 3/3 | 1/3 | 0/3 | 0/3 | ND | 1280, 806, 1810 |
| 1F | H3N2 | 2 days | INF |  | 2/7 | 0/7 | 1/7 | ND | 160, 404, 906, 320, 1280, 906, 320 |
|  |  |  | Naïve | 6/7 | 0/7 | 0/7 | 3/7 | ND | 320, 160, 160, 640, 640, <14, 1280 |
| 1G | H3N2 | 5x 8h | INF |  | 2/3 | 0/3 | 0/3 | ND | 320, 2560, 3620 |
|  |  |  | Naïve | 2/3 | 0/3 | 0/3 | 2/3 | ND | 1810, 1280, <14 |
| 2B | H1N1pdm | 7 days | INF |  | 1/3 | 3/3 | 0/3 | 1613, 1613, 1280 | NA |
|  |  |  | Naïve | 3/3 | 1/3 | 0/3 | 0/3 | 160, 1280, 52 | 806, 226, 905 |
| 2C | H1N1pdm | 2 days | INF |  | 3/3 | 1/3 | 0/3 | 1280, 2560, 1280 | 3620, 1810, 1810 |
|  |  |  | Naïve | 2/3 | 0/3 | 0/3 | 0/3 | 1280, <14, 1280 | 1280, 806, 1810 |
| 3B | H3N2 | 2 days | INF |  | 1/4 | 0/4 | 0/4 | 3620, 3225, 2560, 3620** | 160, 320, 1280, 403 |
|  |  |  | Naïve | 0/4 | 3/4 | 0/4 | 0/4 | 2560, 3620, 1613, 3620** | <14, <14, <14, <14 |
| 3C | H3N2 | 2 days | INF |  | 1/4 | 0/4 | 0/4 | 453, 40, 226, 160 | 906, 640, 906, 906 |
|  |  |  | Naïve | 2/4 | 0/4 | 0/4 | 0/4 | NA | 100, 40, <14, <14 |
| 3D | H3N2 | 2 days | INF |  | 0/3 | 0/3 | 0/3 | NA | 202, 160, 320 |
|  |  |  | Naïve | 0/3 | 0/3 | 0/3 | 0/3 | 640, 80, 453 | <14, <14, <14 |

\* Temperature increase is defined as >1.5deg from day 0 temperature. Weight loss determined as > 10% of day 0 weight
<sup>^</sup> Antibody titers of day 14 are presented based on microneutralization assay. All day 0 sera had a titer <14. Data for animals from Figure 2 present antibody titers from day 14 primary or secondary infection.
ND = Not determined; NA = Not applicable
\*\* From day 21 not day 14

